# Introducing multifactorial electroculturomics: Alternating Current Electric Pulses, combined with mild thermal treatment, exhibit antimicrobial or stimulatory effects on bacterial pathogens and enteroviruses, implying prospects for targeted microbiomics applications

**DOI:** 10.1101/2023.11.24.568545

**Authors:** Grigoria Spanou, Maria Daskou, Manousos E. Kambouris, Chrysanthi Mitsagga, Dimitris Mossialos, Aristea Velegraki, George P. Patrinos, Ioannis Giavasis

## Abstract

**AIMS:** To surrogate chemical and high-energy microbicidals, Electroceuticals may be used as a stand-alone or combined treatment under the guise of Electroculturomics.

**METHODS AND RESULTS:** Using high and low settings of Intensity and Frequency of a medical-rated instrument (TENS) of alternating current the viability and propagation of seven pathogenic bacteria and one enterovirus of environmental and medical importance were tested *in vitro*, in order to establish the interaction of electroceuticals and mild pasteurization protocols and identify potential synergies and/or antagonism of these treatments. The combined regimen showed synergy, following the prerogatives of the Bioelectric Effect, and antagonism. High frequency (800Hz) rather than low (2 Hz) seems detrimental, while intensity (10 or 1 mA) seems almost inconsequential, while longer sessions enhance detrimental effects but short exposure may be beneficial.

**CONCLUSIONS:** No single treatment seems optimal for all tested bacteria. High frequency can be effective against low titers of Enterovirus, but at higher titers, the effect may be reversed. Case-specific effects on microbial growth patterns seem to be the norm.

**SIGNIFICANCE AND IMPACT OF STUDY:** Diverse mechanisms of microbicidal or stimulatory activity are implied, allowing individualized uses and targeted applications in food and environmental safety, therapeutics and industrial bioprocessing.

## INTRODUCTION

The extended use of antibiotics against infectious diseases, zoonoses as well as for various agricultural and environmental purposes has been in the verge of abuse (English and Gaur, 2010; Kleinkauf and Monnet, 2013). This tendency reinforced the natural resistance mechanisms of microbes, resulting in low efficacy and environmental burden. Similarly, synthetic preservatives have been extensively used for the preservation of perishable food products for decades, but consumers’ concerns for their potentially health-compromising effects and the modern market preference for more natural and low-processed foods has fueled the search for natural antimicrobials and mild antimicrobial treatments (Skenderidis *et al*., 2019; Nair *et al*., 2020; Didaras *et al*., 2021). Therefore, the need for novel, better and more affordable amenities has incited research for alternative approaches. Electrostimulation (ES) in different forms and shapes, occasionally termed collectively as Electroceuticals (Kambouris *et al*., 2014) in a broader sense, promises cost-effective, adaptable and flexible solutions either complimentarily or alternatively to other approaches (Giladi *et al*., 2008; Kambouris *et al*., 2014, 2017).

However, the precise role of ES in microbial growth has not been yet determined (Hristov and Perez, 2011; Siamoglou *et al*., 2020). Electrical modalities have been tested for more than a century to regulate microbial growth in order to induce specific growth routines but also to decontaminate objects (Kermanshahi and Sailani, 2005; Valle *et al*., 2007; Petrofsky *et al*., 2008). An extensive, though not cohesive, body of research has been conducted with vastly different cultures, organisms, equipment and objectives and it clearly substantiates the potential of such an approach (Kambouris *et al*., 2014, 2017; Stathoulias *et al*., 2020). Conductive amenities, where electric current is directed through the cells/organisms, result in a targeted, efficient treatment with low residual energy to the environment. Ohmic effects, both to the target and its environment, including but not limited to growth substrates (Davis *et al*., 1992; Kambouris *et al*., 2014; Siamoglou *et al*., 2020; Stathoulias *et al*., 2020), offer advantageous conditions for suppressive purposes, thus suggesting synergy with thermal amenities. Exposing cells and organisms to electric current for time, charge and intensity values below thresholds of ohmic damage can be beneficial for the microbe in stimulating growth routines (Kambouris *et al*., 2014, 2017; Stathoulias *et al*., 2020), such as multiplication, expansion, primary and secondary metabolite production, and self-healing; the latter is of vital importance in infection control and in sterilization, as it may undermine the effect of other amenities used in parallel or in tandem. Treatment of microbial cultures with conductive amenities causes low electromagnetic pollution compared to respective fields and microwaves while allowing manipulation of the growth of different microbiota, especially during propagation, as dividing cells are more susceptible to ES than growing, stable or static cells (Siamoglou *et al*., 2020; Stathoulias *et al*., 2020). Previous experience shows that the results of ES on microbial growth may be highly diverse by microbe and not following linear correlations (Stathoulias *et al*., 2020). The ability to affect the growth of microbes, in either dynamic or kinetic terms, may offer operational options beyond the chemical and genetic regulation of microbes for agro-industrial, food and pharmaceutical applications.

## MATERIALS AND METHODS

### Equipment

The portable device, Mio-Ionotens (I-Tech Medical Division, Martellago Italy) was used for the experiment. Mio-Ionotens is a portable and battery-powered device that generates Transcutaneous Electrical Nerve Stimulation (**TENS**) and ionophoresis currents. It is a device of two independent channels, providing either high or low power at its outputs and performing a series of TENS and iontophoresis programs. Each channel may treat one or two targets; in the second case the use of the provided splitter cabling is required, thus dropping the intensity output to half the nominal value.

### Cultures of microorganisms

Different pure cultures of common bacterial pathogens (both Gram positive and Gram negative ones) implicated in foodborne disease and a Poliovirus type Enterovirus of intestinal/environmental/food origin that may cause potentially fatal disease (Pallancsh, 2013) were used as targets of antimicrobial activity in this study.

In order to prepare fresh stock cultures of bacteria, 1ml of previously prepared stock culture was inoculated into 10ml Tryptone Soy Broth (TSB) tubes. These were incubated at the appropriate temperature and time for each microorganism. *Escherichia coli* <u>NCTC 9001</u> was incubated at 37 °C for 24 h, *<u>Salmonella enterica</u>* <u>subsp. *enterica serovar* typhimurium NCTC 12023</u> (there after called *Salmonella typhimurium*) at 37°C for 48h, *Campylobacter jejuni* was incubated under microaerophilic conditions (by addition of 1 mL of sterile paraffin) at 37°C for 48h, *Staphylococcus aureus* <u>NCTC 6571</u> at 37°C for 48h, *Listeria monocytogenes* NCTC 7973 at 37°C for 48h, *Bacillus cereus* <u>NCTC 7464</u> at 37°C for 48h and *Clostridium perfringens* <u>NCTC 13170</u> at 37°C for 48h (all cultures were obtained from NCTC culture collections, UK).

With regard to the Poliovirus type 1 enterovirus, the strain Sabin (LSc, 2ab) from the American Type Culture Collection (ATCC, Rockville, Maryland, USA) with accession number V01150 was used in this study. The titer of the virus was calculated as means of Median Tissue Culture Infectious Dose /TCID_50_ (Hierholzer and Killington, 1996) after the inoculation of the virus in Rhabdomyosarcoma (Rd) cells (human rhabdomyosarcoma cell line—CCL-136™, ATTC). Subsequently, serial dilutions were performed in order to obtain the desirable titers of 10^6^ and 10^2^ TCID_50_/0.1ml.

### Treatment of microorganisms

For treatment of bacteria with the mio-IONOTENS device (Fig.1), the electrodes of one channel of the device were inserted aseptically in the culture vessels (glass test tubes) containing 5 ml fresh Tryptone Soy Broth (TSB, Neogen, USA). Two different ES intensities were used, 1mA and 10mA (low and high intensity), and two different frequencies, 2Hz and 800Hz (low and high frequency), resulting in 4 treatment cases for 1 minute, 10 minutes and 30 minutes, resulting in a total of 12 electrostimulation treatments per strain. These four ES programs were used in a stand-alone context (1^st^ experiment), and also combined with a standard thermal pasteurization step (2^nd^ and 3^rd^ experiments), using a LabTech water bath (Tecotec, Vietnam) set at 65°C. This relatively mild pasteurization temperature was chosen in order to test the viability of pathogens in a context of minimal processing. Two combinations of ES-pasteurization were tried, the *tandem*, where the pasteurization step was implemented after the electrostimulation treatment (2^nd^ experiment), and the *parallel*, where ES and pasteurization were implemented simultaneously (3^rd^ experiment). The alpharithmetic coding of each treatment regimen is shown in Table 1. The order applies to all three batches of treatments, referring to treatment time, in all tested bacteria (Figure 2). Each alpharithmetic code refers to 21 physical samples (seven microorganisms times three treatment durations).

**Table 1:** Sample codes by treatment conditions. (irrespective of treatment time and bacterial species)

| Sample code | Treatment regimen | Experimental group |
| --- | --- | --- |
| 1 | Untreated control | ES only<br>(1<sup>st</sup> experiment) |
| 2E | 800Hz,10mA |  |
| 3E | 2Hz,10mA |  |
| 4E | 800Hz,1mA |  |
| 5E | 2Hz,1mA |  |
| 1P | Control (pasteurized, no ES) | ES followed by pasteurization<br>(2<sup>nd</sup> experiment) |
| 2E+P | 800Hz,10mA |  |
| 3E+P | 2Hz,10mA |  |
| 4E+P | 800Hz,1mA |  |
| 5E+P | 2Hz,1mA |  |
| 1P | Control (pasteurized, no ES) | ES during pasteurization<br>(3<sup>rd</sup> experiment) |
| 2EP | 800Hz,10mA |  |
| 3EP | 2Hz,10mA |  |
| 4EP | 800Hz,1mA |  |
| 5EP | 2Hz,1mA |  |

For the electrostimulation of the enterovirus culture, Sabin 1 reference strain in two different concentrations (populations) was subjected in the first four ES treatments described in Table 1. Namely, the Enterovirus samples were subjected to: 800Hz x 10mA (Sample 2E), 2Hz x 10mA (Sample 3E), 800Hz x 1mA (Sample 4E), and 2Hz x 1mA (Sample 5E). A control with no treatment (Sample 1) was also used for comparison. These four ES treatments were also combined with simultaneous heating at 60^ο^C for 30 min only, in an effort to test for synergistic effects of electrostimulation and pasteurization on the desirable reduction of the virus titer. Pasteurization alone at 65^ο^C would eliminate the titer of 10^2^ TCID_50_/0.1ml and reduce the titer of 10^6^ TCID_50_/0.1ml to barely detectable levels, thus making synergies undetectable if not irrelevant. Thus, a milder pasteurization temperature (60^ο^C) was used for the Sabin strain of Poliovirus type 1, combined-or not with electrostimulation.

### Enumeration of Bacterial Population

After the end of the treatment, the population of each microorganism was assessed by viable counts of colonies on selective solid (agar) media. *E. coli* was grown on Chromogenic Tryptone Bile X-glucuronide (TBX) Agar (Neogen. USA) at 37°C for 24h; *Salmonella typhimurium* on Xylose Lysine Desoxycholate (XLD) Agar (Merck, USA) at 37°C for 24h; *Campylobacter jejuni* on Campylobacter Blood-Free Selective Agar Base with Campylobacter selective supplement (Neogen, USA) under microaerophilic conditions in anaerobic jars with Anaerocult C (Merck, USA) at 37°C for 72 h; *S. aureus* on Baird Parker Agar Base (Neogen, USA) with egg yolk tellurite at 37°C for 48h; *L. monocytogenes* on Harlequin® Listeria Chromogenic (ALOA) Agar with L. monocytogenes selective supplement (Neogen. USA) after incubation at 37 °C for 48 h; *B. cereus* (vegetative cells) on Bacillus Cereus Agar Base with Polymixin B selective supplement (Neogen, USA) at 37°C for 48h; and *C. perfringens* (vegetative cells) on TSC Perfingens (Tryptose Sulfite Cycloserine) Agar Base + D-cycloserine selective supplement (Neogen, USA) at 37°C for 48h in anaerobic jars with Anaerocult A (Merck, USA). Prior to enumeration, each liquid culture in TSB was properly diluted in serial dilutions with Maximum Recovery Diluent (MRD). Three serial dilutions were used for enumeration of each microorganism and were tested in accordance with the standard criteria of microbiological enumeration methods of American Society of Testing and Materials (ASTM, 1998). The reported values represent the means of accepted values of the three serial dilutions of each series of experiment. A series is defined by one control (no treatment) and 4 different ES treatments as described in Table 2.

**Table 2.** Real-time PCR results presented as Ct values after electrostimulation processing.

|  | 1 | 2E | 3E | 4E | 5E |
| --- | --- | --- | --- | --- | --- |
| **S1 10<sup>2</sup> TCID<sub>50</sub>** | 18.06 | 24.52 | 16.18 | 21.54 | 18.27 |
| **S1 10<sup>6</sup> TCID<sub>50</sub>** | 16.44 | 17.48 | 17.76 | 17.77 | 17.43 |

**Table 3.** Real-time PCR results presented as Ct values after electrostimulation processing combined with heating in 60°C.

|  | 1 | 1P | 2E+P | 3E+P | 4E+P | 5E+P |
| --- | --- | --- | --- | --- | --- | --- |
| **S1 10<sup>2</sup> TCID<sub>50</sub>** | 18.89 | 29.97 | No Ct | No Ct | No Ct | 37.15 |
| **S1 10<sup>6</sup> TCID<sub>50</sub>** | 16.29 | 25.59 | 27.91 | 16.74 | 22.52 | 15.8 |

### Cell culture of the treated viral samples

After the treatment of the viral strain in concentrations of 10^2^ and 10^6^ TCID_50_/0.1ml., 200μl of each sample were inoculated in Rd cells, which were seeded in cell culture tubes and grown in D-MEM (Sigma-Aldrich, St Louis, MO, USA) supplemented with 2% fetal bovine serum (Sigma-Aldrich, St Louis, MO, USA) 24h before the inoculation. Then, the culture tubes were incubated at 37^ο^C and tested daily for cytopathic effect (CPE) development. Non-treated Sabin 1 samples in concentrations of 10^2^ and 10^6^ TCID_50_/0.1ml in Rd cells were used as positive virus controls in order to differentiate CPE due to virus activity. In addition, non-infected Rd cells were used as negative control.

### Viral RNA extraction and Reverse Transcription

The viral RNA was extracted from the samples by using the guanidine thiocyanate (GuSCN) extraction protocol (Casas et al., 1996). At the final step the pellet was dried and dissolved in 100μl of DNase-RNase free double-distilled sterile water (DEMO S.A, Athens, Greece).

Subsequently, the extracted RNA from all tested samples was subjected to a Reverse Transcription reaction using random primers d(N7) (Macrogen, South Korea) and FastGene Scriptase II (Nippon Genetics, Japan). The reaction was divided in two steps. For the first step the reaction mixture consisted of 5 pmol of random d(N7) primers (Macrogen, South Korea), 2 mM of dNTPs (Invitrogen Life Technologies, Paisley, UK) and RNase-free distilled water up to a volume of 7 μl. Then 5 μl of the extracted viral RNA with 7 μl of the first reaction mixture were incubated for 5 min at 65^ο^C. Then the samples were rapidly transferred on ice and 8 μl of a second reaction mixture consisted of 1X FastGene Buffer, 0.01 mM of DTT, 20 units of ribonuclease inhibitor, 100 units of FastGene Scriptase II (Nippon Genetics, Japan) and RNase-free distilled water up to a volume of 8 μl, were added in each sample and incubated at 25^ο^C for 10 min, 42 for 50 min and 70^ο^ C for 15 min.

### Real-Time PCR assay of viral samples

The reduction of the virus titer was assessed using a Real-Time PCR assay. The Ct (Cycle threshold) value of treated samples was compared to those of non-treated virus controls in order to evaluate the effectiveness of electrostimulation as a stand-aloneorpart of combined regimen with thermal processing, in the reduction of the virus titer. Ct levels are inversely proportional to the amount of target RNA in the sample; as a result the lower the Ct value, the greater the amount of the virus in the sample.

For this reason, 3μl of cDNAs were added in a 0.2ml tube containing 17μl of the reaction mixture. The mixture included 1X FastGene Mix (Nippon Genetics, Japan), 50nM ROX Low, 10pmol of each universal enterovirus primer EV2 (5’- CCCCTGAATGCGGCTAATC-3’) / EV1 (5’-GATTGTCACCATAAGCAGC-3’) targeting the 5’-UTR region, which is the most conserved region among *Enterovirus* genus and used widely for their detection (Monpoeho *et al*., 2000) and ddH_2_O up to 20μl. The reaction was conducted on the Mx3005P^@^ Real Time PCR instrument in the following cycling conditions: 95^ο^C for 2 minutes, 40 cycles of 95^ο^C for 5 seconds and 60^ο^C for 30 seconds, followed by a melting curve analysis step.

## RESULTS

The results of the ES treatment of bacterial pathogens, with or without thermal pasteurization, are resented in Figure 2 (a-g) per species. The similar values of two “1P” samples (pasteurized samples of two separate experiments) in all strains and treatment times show a coherent, reproducible growth and serve as quality control for the accuracy and reproducibility of the processing and enumeration methods used.

It becomes apparent from the results of different ES treatments, with or without pasteurization of different pathogenic bacteria, that the effectiveness of ES treatment depends mainly on the duration of treatment, the frequency and only to some extend on the intensity of ES. Furthermore, external factors of importance are the combination with pasteurization (performed synchronously or asynchronously to ES) and the target microorganism.

*B. cereus* vegetative cells seem to be largely unaffected by treatment with ES alone, especially after 1 or 10 min of ES, but show a synergistic effect of ES of 800Hz and thermal processing after 30 min of combined (consequent) treatment (Fig 2a). Moreover, the simultaneous ES treatment with pasteurization is not only synergistic, but practically eliminates all vegetative cells. For *C. jejuni*, this synergistic effect between ES and pasteurization is also observed when the treatment lasts for at least 10 min and is more obvious at 30 min synchronous regimen (Fig. 2b). *C. perfringens* vegetative cells are strongly inactivated with asynchronous combined treatment but even more so with synchronous (Fig. 2c). For *E. coli*, apart from the synergism between pasteurization and ES treatment, which is particularly lethal when synchronous, there is a significant microbicidal effect caused by ES treatment alone, if applied for 30 min (Fig. 2d). In the case of *L. monocytogenes* the ES of 800Hz x 1mA for either 10 or 30 min is highly lethal when asynchronous to pasteurization, while synchronous regimen drops viable cells below detectable levels. This contrasts interestingly with sole pasteurization (without ES) scores, whence high population of viable cells can be detected, at ∼4 log cfu/ml (Fig. 2e). With *S. typhimurium* synergism of ES and pasteurization is detectable even if for only 1 min of treatment and, in contrast to most other tested bacteria, the susceptibility window seems to appear at 2Hz ES (Fig. 2f). For *S. aureus* the ES treatment is potentially destructive by itself, but may also lead to better survival during pasteurization, especially if ES is applied for a very short time (1 min); a combined pasteurization and ES treatment shows a synergistic lethal effect if applied for longer times, especially in synchronous formats (Fig. 2g).

Overall, the simultaneous application of pasteurization and ES offered the best microbicidal results, which, depending on the species, where up to 5 orders of magnitude, as in the case of *E. coli* when treated for ten or thirty minutes with a synchronous combined regimen of ES (irrespective of the ES specifics) and pasteurization (Fig. 2d). It is also evident that different species react differently to the tested regimens in both absolute and relative terms. *S. typhimurium* seems least affected and *C. perfringens* most affected by ES alone; the latter, subjected to ES only for 1min at 2Hz, 10mA or at 800Hz, 1mA showed 2 orders of magnitude lower viable counts than the untreated control.

Even more interesting is the indication for potentially stimulatory effect of ES or ES- heat treatment, which seems to affect beneficially the bacterial growth or survival during pasteurization, as in the case of a short, 1 min ES treatment of *B. cereus*, *C*. *jejuni* and *S. aureus*. Even when combined with pasteurization the results of ES are not always detrimental: *C. jejuni* when treated with ES at 800Hz, 10mA and 2Hz, 1mA under pasteurization for 1min showed higher viable counts than the pasteurized control. These results indicate a healing/repair or otherwise protective mechanism of the specific ES regimen against the detrimental effect of pasteurization.

Similarly, different ES settings may show similar results in a series, or wildly different. The latter is the case of synchronous regimens applied to *S. aureus* for 30min: a frequency of 800Hz produces more intense detrimental effect than one of 2Hz, irrespective of the intensity. In the former case *E. coli* under all synchronous regimens suffers an almost lethal, not simply detrimental effect of 5 orders of magnitude.

The results of Real-Time PCR assay of the Enterovirus samples depicted in Table 2 showed that electrostimulation processing at 800Hz, regardless of intensity (samples 2E and 4E), reduced the virus titer of treated Sabin 1 samples when the initial viral concentration was 10^2^ TCID_50_/0.1ml. In contrast, in sample 3E the virus titer had actually risen, which indicates a beneficial/supportive effect on viral propagation. In sample 5E (2Hz x 1mA) the titer of the virus remained practically unaffected. When the initial viral concentration was high, at 10^6^ TCID_50_, the titer remained unaffected by any of the ES regimens used.

Interestingly, when the ES protocols were combined with heating at 60^ο^C a synergistic detrimental effect was observed in Sabin 1 titer in both viral concentrations that were used; the antiviral effect was significantly increased compared to similar ES-only treatments, as presented in Table 3.

For the low viral concentration (10^2^ TCID_50_) the antiviral effect of ES treatments 2E, 3E, 4E combined with pasteurization was so pronounced that no viral genome was detected after treatment (“No Ct”). It became detectable only when pasteurization was combined with the mildest ES regimen of E5 (2Hz x 1mA) with a Ct value of 37.15 (Table 3). These results indicate a significantly increased reduction of the titer of Sabin 1 when starting from 10^2^ TCID_50_/0.1ml in all ES-pasteurized samples, compared with the reduction achieved with sole pasteurization.

Having said that, the absence of measurable Ct values does not necessarily mean complete elimination of the virus, but only a relative reduction of the viral titer, which might leave active virions that might multiply eventually in subsequent passages of cultures in Rd cells. Nevertheless, even the functional neutralization of Enteroviruses in water and food by ES would be of great importance.

As already mentioned, when starting from a high viral concentration of 10^6^ TCID_50_/0.1ml, the titer of Sabin 1 was synergistically decaying due to the combined ES and pasteurization only in sample 2E+P (ES of 800Hz, 10mA), while all other ES treatments seemed to alleviate the antiviral effect of pasteurization. This indicates that the concentration of the virus is a decisive factor that affects the performance of ES treatment. All ES treatment regimens seemed to enhance the antiviral effect of pasteurization when a low titer was used, but only the most intense ES treatment of high frequency-high intensity (800 Hz x 10 mA) was capable of exerting an additional microbicidal effect upon pasteurization, when a high virus titer was used. The reason for an actually neutralizing effect to the pasteurization impact exerted by the other three ES treatment protocols remains unfathomable. It can be hypothesized, though, that in the case of high viral titer the vibrating effect of electrostimulation is attenuated and may actually facilitate (instead of prevent) the attachment of virions to host cells.

## DISCUSSION

The sector of electroregulation in microbiota has received considerable attention since 1932 (Tracy, 1932) especially in the field of food science (Berovic et al, 2008; Hristov and Perez, 2011); mostly for suppressive applications, as is decontamination and conservation (Tracy, 1932; Ranalli *et al*., 2002), but for inductive applications also, for achieving higher yields in foodstuff and beverage microbiotic maturation and for more efficient growth of microbial cultures for such use. The technology is not expensive, nor overly sophisticated; the main issue has always been a lack in cohesive experimental planning with clearly defined targets and research strategy.

This project resides in culturomics and vies for a compact, standardized and cohesive format with a very limited scope of tunable parameters to explore effects on microbial growth so as to identify possible prospects for future development of application-specific protocols and basic research. The use of off-the-self equipment, methods and technology to establish an innovative approach in manipulation of microbial growth curtails developmental risks and time, and thus focuses on the translational, commercial dimension of respective research with some compromising of the purely scientific aspects. Still, preliminary research is well tolerant of such compromises.

The stimulus has been a previous study with fungal solid cultures that detected with some certainty both detrimental and beneficial effects of the same regimen, depending on species/strain and dosage (Stathoulias *et al*., 2020). This is somewhat contradicting the conventional wisdom of the Bioelectric Effect principle, which produces synergy between electromagnetic treatment schemes and standard antimicrobial approaches, especially (bio)chemical in origin (Costerton *et al*., 1994; Giladi *et al*., 2008), but not something utterly unexpected (Kambouris *et al*., 2017), as infections have been steadily considered counter-indication for the prescription of electroceuticals, despite available preliminary evidence (Kambouris *et al*., 2014).

In this study a different procedure was followed, with extreme conditions (high and low ES intensity as well as frequency) tested in bacterial liquid cultures and viral cell cultures with a clear intent to identify a regimen for improved antimicrobial treatment of sensitive liquids as found in the food industry (e.g. potable water, fresh juices, etc.) or in environmental samples (such as sewage water). ES seems to represent a novel and potentially powerful tool for the deactivation of pathogens in environmental and food matrices, especially when combined with mild thermal processing, which is significant to food preservation and safety and can allow for savings in energy costs and carbon emissions, as opposed to high-temperature pasteurization. In the case of Enteroviruses, which are not only abundant in sewage water and feces, but also resistant to chemical treatment/deactivation due to the lack of envelope (Kyriakopoulou *et al*., 2010; Iritani *et al*., 2014), it becomes particularly crucial to develop effective protocols for their inactivation.

Still, another priority of this work has been to identify potential beneficial effects for improved cultivability and enhanced growth (or survival under stress conditions) of economically important cultures, implicated in the production of important metabolites in fermented foods or industrial fermentations, where an improved growth rate is of cardinal importance (Bouki *et al*., 2020; Schmidt, 2005). The concept of this study prioritized highly the testing of different ES combinations and the exhibited suppressive or stimulatory effects towards different microorganisms.

The biological activity - and its underlying mechanisms - are all somewhat hazy at the moment, but the diverse effects caused by the same modalities onto different organisms show that a complex network of interactions should be considered at play, rather than a specific, linear pathway and a defined set of implicated genes and procedures. One may only speculate on the effects of different ES regimens on different growth parameters. Such are, indicatively and not exhaustively: The lag or acceleration of the onset of the exponential phase, which is of major importance in curtailing contamination and infection. The balancing of biomass increase versus (radial) expansion growth, which is important in infection control and in biomass production. The balance in producing primary and secondary metabolites for industrial application, such as – but not restricted to - food and beverage industry (Brian, 1951; Perez *et al*., 2007; Detman *et al*., 2019) or catabolizing substrates in biodegradation/bioremediation applications (Kormas and Lymperopoulou, 2013). The resistance profiles to antibiotics, antiseptics, synthetic preservatives or natural antimicrobials and other antimicrobial amenities, especially through the bioelectric effect (Costerton *et al*., 1994) that increases the microbicidal potential of antimicrobial treatments, decontamination protocols (Davis *et al*., 1991) and therapeutics (Carley and Wainapel, 1985; Spadaro, Chase and Webster, 1986; Ramadhinara and Poulas, 2013; Kambouris *et al*., 2014). Our results indicate that an individualized treatment of specific pathogens under specific ES conditions might be possible, without exposing other (beneficial) members of a microbiome to intense and destructive treatment. This would be especially useful in matrices like fermented foods where pathogens or spoilage organisms need to be restricted, while starter cultures and autochthonous microbiota taking part in the fermentation needs to be unaffected, or, if possible, enhanced (Bouki *et al*., 2020). But the principle applies to the human microbiome in different biocompartments, especially when corrupted by aggressive, potentially pathogenic strains or opportunists.

The results with Enteroviruses also hint that electrostimulation combined with heating provides better results in the reduction of the virus titer and inhibition of viral growth, especially at high frequency-high intensity settings of ES, but this effect depends on the concentration of the virus as well. Under high viral titer and low frequency (2 Ηz) ES the opposite effect may occur. Therefore, this electrostimulation method could potentially limit Enteroviruses, if at low concentrations, in environmental and food matrices (e.g. water, seafood, fresh produce). It could also apply to other virus families and evolve to a new preventive or therapeutic approach against common viral pathogens. On the other hand, one cannot exclude a potential stimulatory effect that low frequency ES may exert on high titers of Enteroviruses or other viruses), which could also be useful in quite some applications. Such are: The enhanced production of viruses to use as vectors for vaccines. The increase of the efficiency of vaccines using viral vectors, to speed up the development of immunity after vaccination. The more efficient use of viral/phagal vectors for transfection in biotechnological applications and research. And, last but not least, the increase of the efficiency of bacteriophages and virophages for food/animal decontamination (Zalewska-Piątek and Piątek, 2021) or when used as biotherapeutics.

The use of electrical modalities, especially low intensity (and low power) electrostimulation, is a promising, green alternative to the environmentally heavy footprints of chemical and intense thermal processing for food preservation, environmental decontamination and therapeutics (Cavicchioli *et al*., 2019). Besides, the global consumer and therapeutic trends point to minimizing the use of synthetic food preservatives, antibiotics and optimizing the nutritional value of food by mild or minimal processing (Fardet, 2016). It may also alleviate the need for dedicated, genetically engineered microbiota for specific biotechnological needs that seem to become a trend in some industrial bioprocesses (Ford and Silver, 2015; Janssen and Stucki, 2020; Schipp *et al*., 2020). Thus, the need for expensive and patented strains might be reduced, if natural ones could be physically induced to increase their efficiency and/or adaptability (Marrero *et al*., 2015; Donati, Castro and Urbieta, 2016). Except for lower costs, a reduced number of engineered strains would result in less genetic pollution and unpredictable recombination in the environment (Schmidt, 2010; Acevedo-Rocha and Budisa, 2016).

Such prospects for achieving targeted manipulation of microbial growth via electrostimulation insinuate a degree of optimization. Since the impact of ES treatment depends heavily on individual cellular characteristics, indiscriminate destruction of the microbiome (e.g. as in the case of antibiotics or chemical disinfectants) will be averted and microbial diversity can be preserved, either in a fermented food matrix, or in the context of the microbiome of diverse biocompartmens, such as the gut. The diverse cell structure characterizing different microbial taxa may be responsible for the highly variable responses to electrostimulation. For instance, highly charged or polarized macromolecules that interact with electric pulses may differ in distribution and quantity, thus resulting in different cell responses to a given ES stimulus. This work aspires to contribute to revealing the unexplored field of microbial electrostimulation as studied and potentially developed through electroculturomics and pave the way for the exploitation of Microbiome-targeting Electroceuticals in Food and Environmental Safety, Industrial Bioprocesses and Human Health applications.

## Authors contribution statement

**Grigoria Spanou** and **Chrysanthi Mitsagga** performed the experiments implicating bacteria, assisted to the optimization of the method for liquid bacterial cultures combined with pasteurization and in the interpretation of the results, and participated in finalizing the manuscript.

**Maria Daskou** performed the experiments implicating viruses, assisted to the optimization of the method for application in viral context and in the interpretation of the results, and in finalizing the manuscript.

**Manousos E. Kambouris** conceived the electroculturomics concept and the generic application of the methodology herein, assisted in setting the specifications for the experimental setup and participated in the interpretation of the results and the preparation of the manuscript.

**Aristea Velegraki** and **George P. Patrinos** assisted in the development of the Electroculturomics concept, the experimental setup and the interpretation and presentation of the results and drafting the Discussion part of the manuscript.

**Dimitris Mossialos** developed the viral dimension of the work, designed the respective experiments, provided their interpretation and with M Daskou authored the part of the manuscript that refers to viruses.

**Ioannis Giavasis** developed the bacterial dimension of the work by adapting the methodology to liquid cultures, designed the respective experiments, provided their interpretation and assisted in the drafting of the manuscript.

---

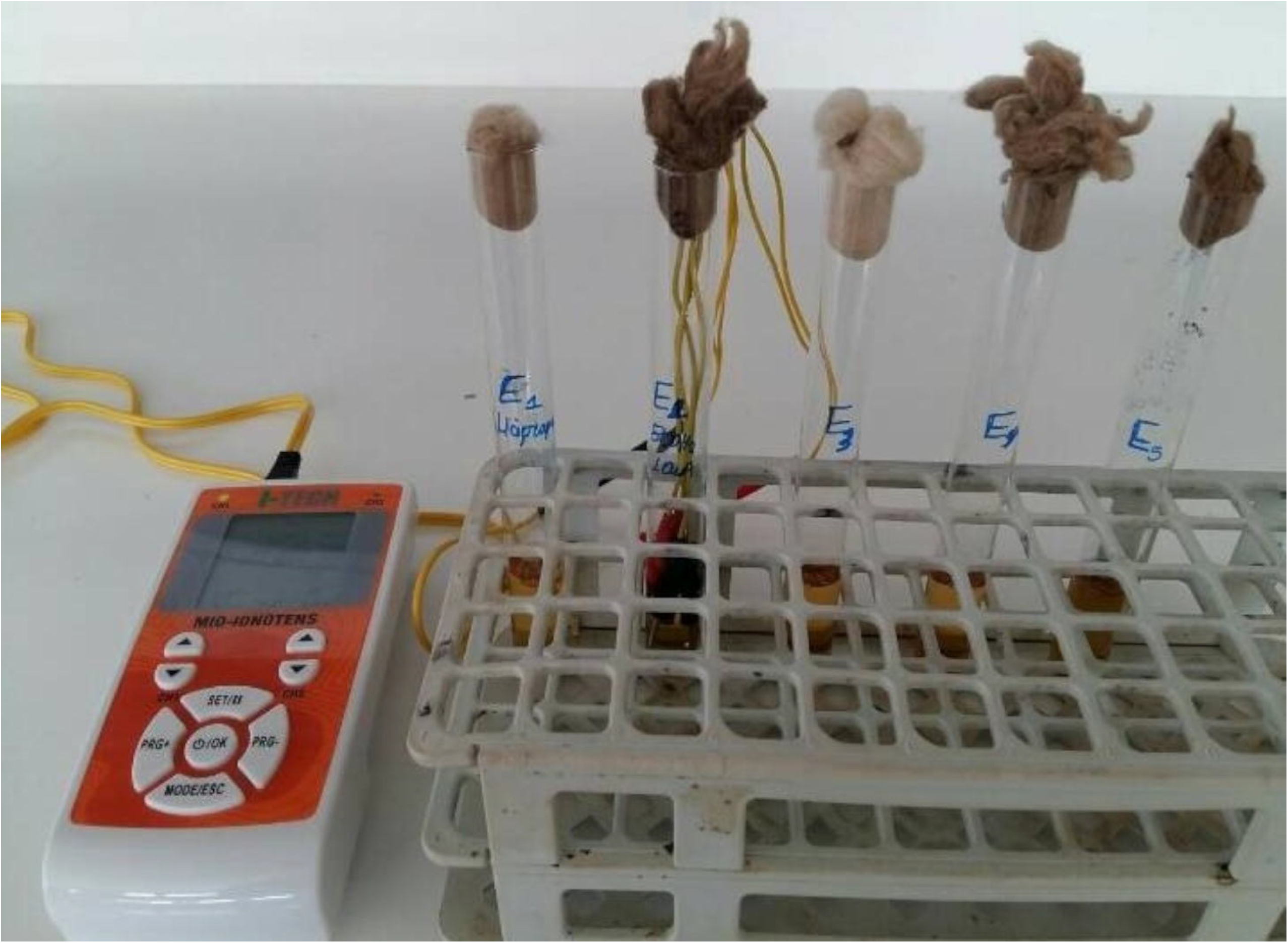

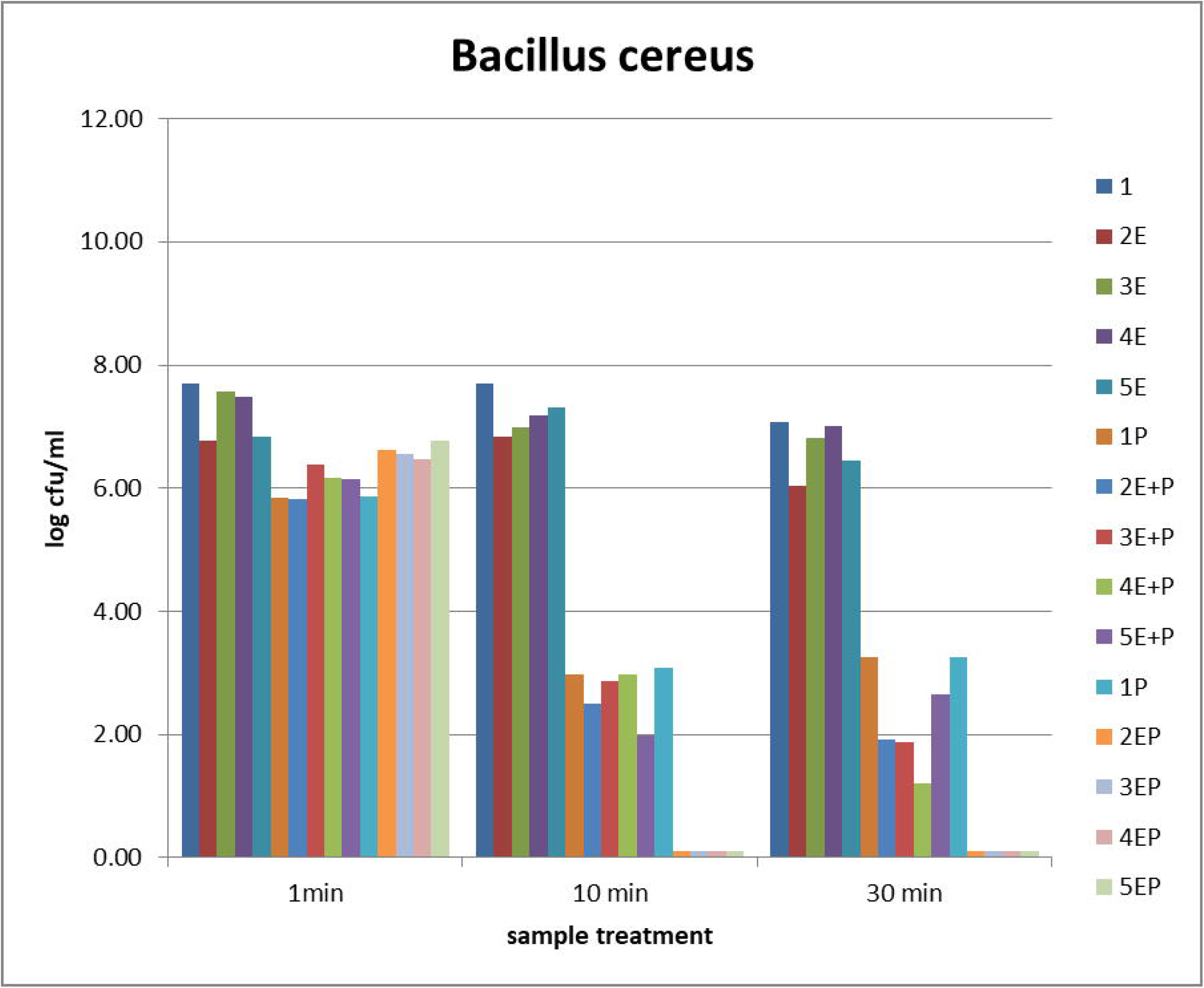
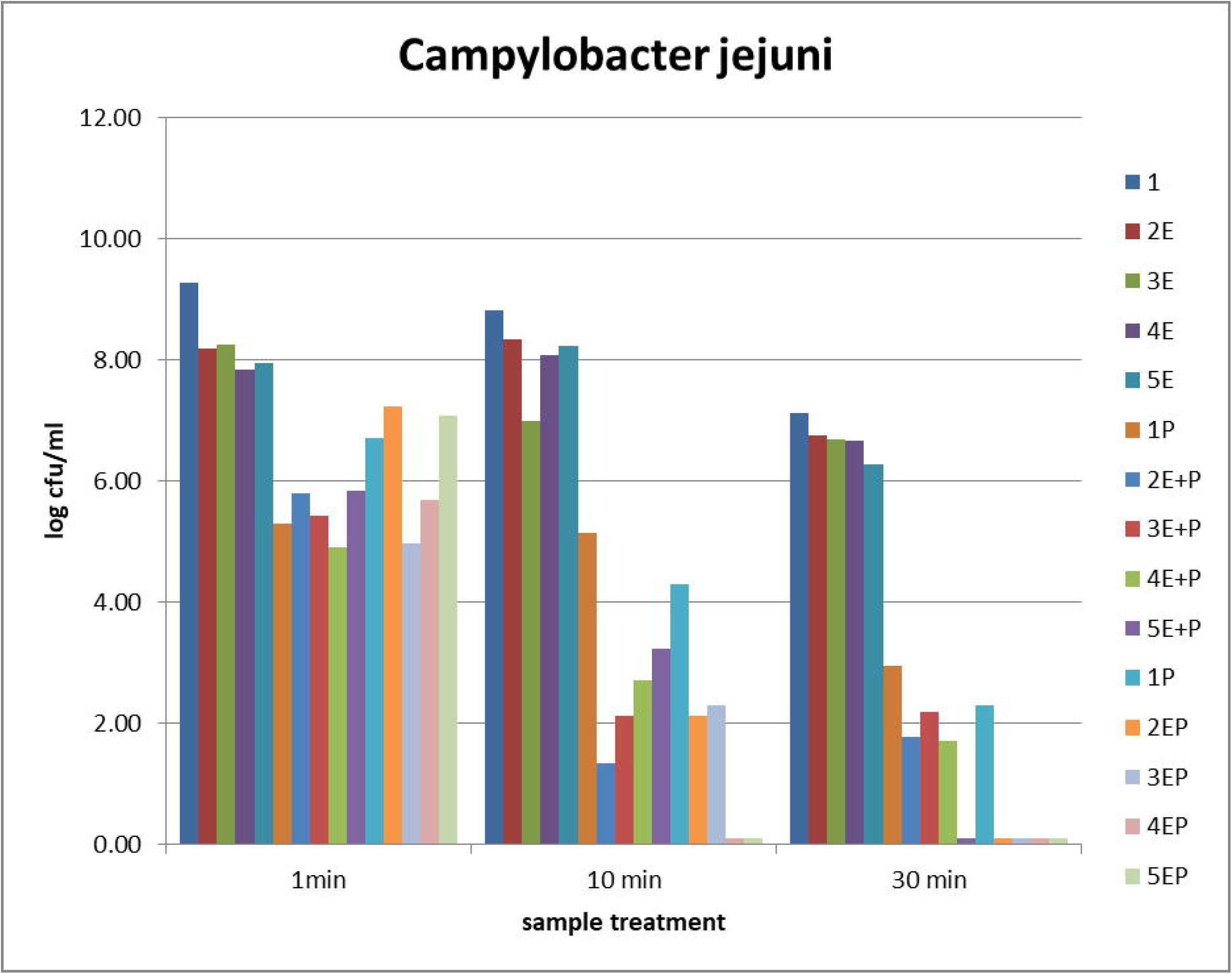
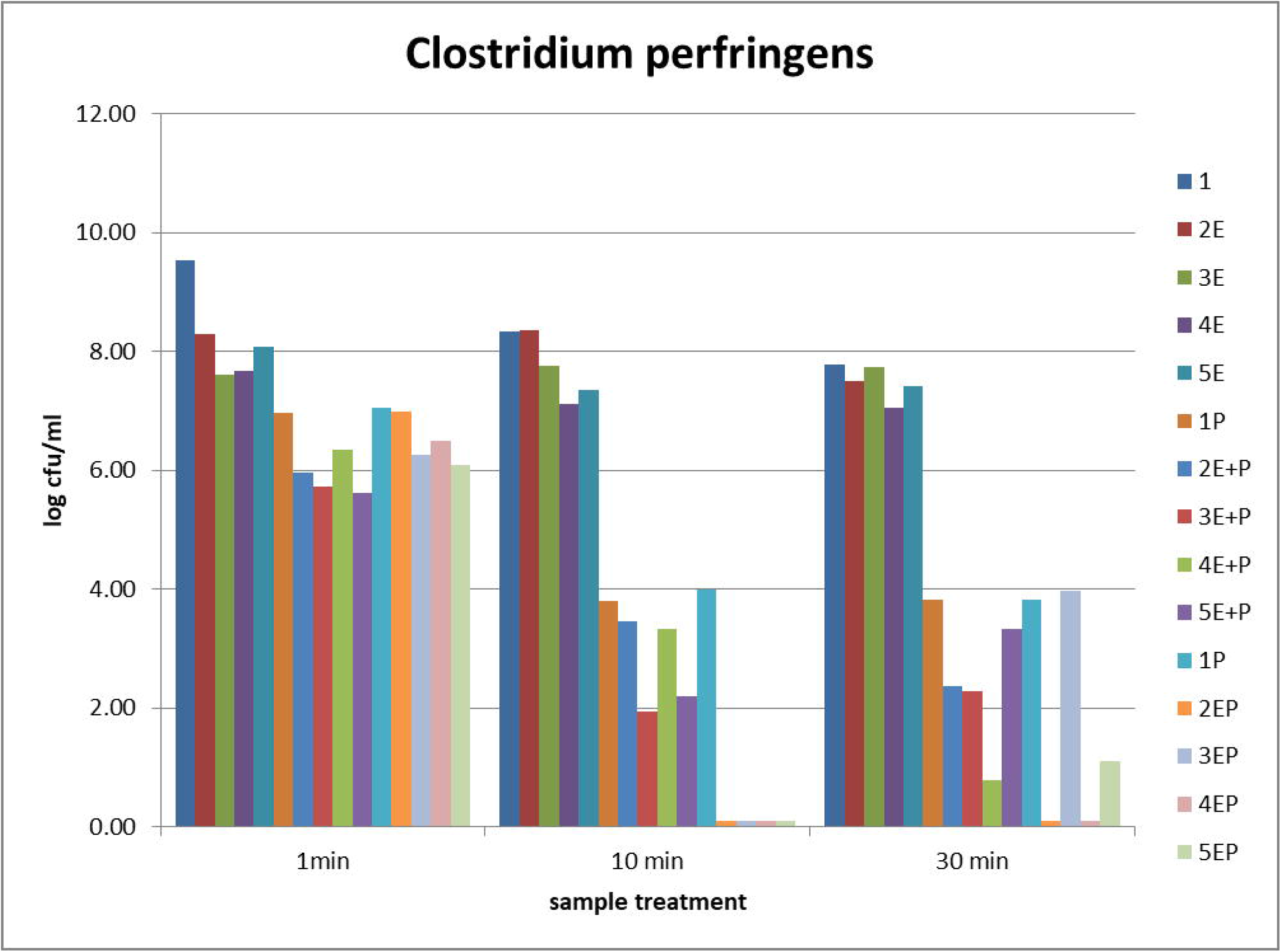
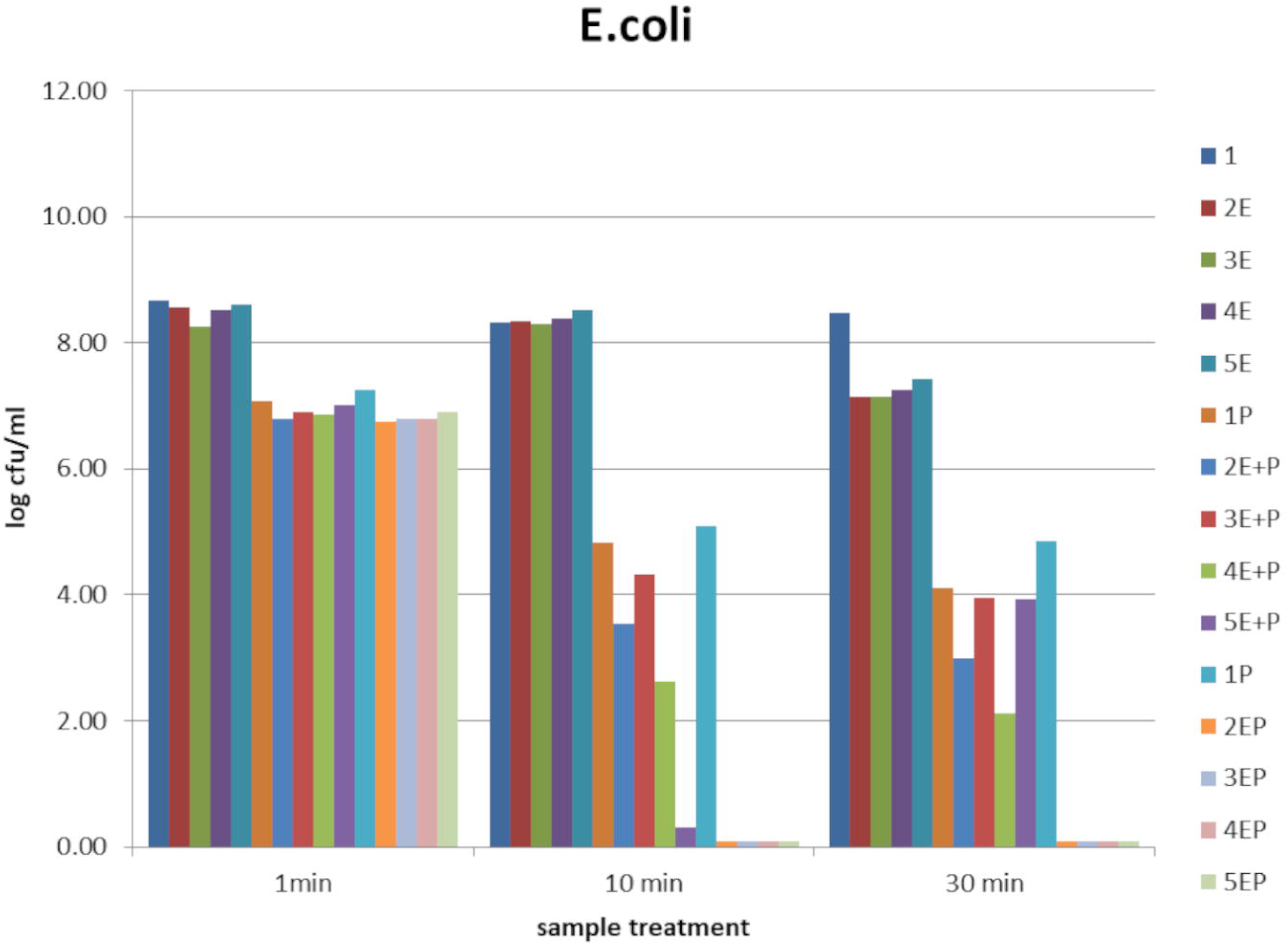
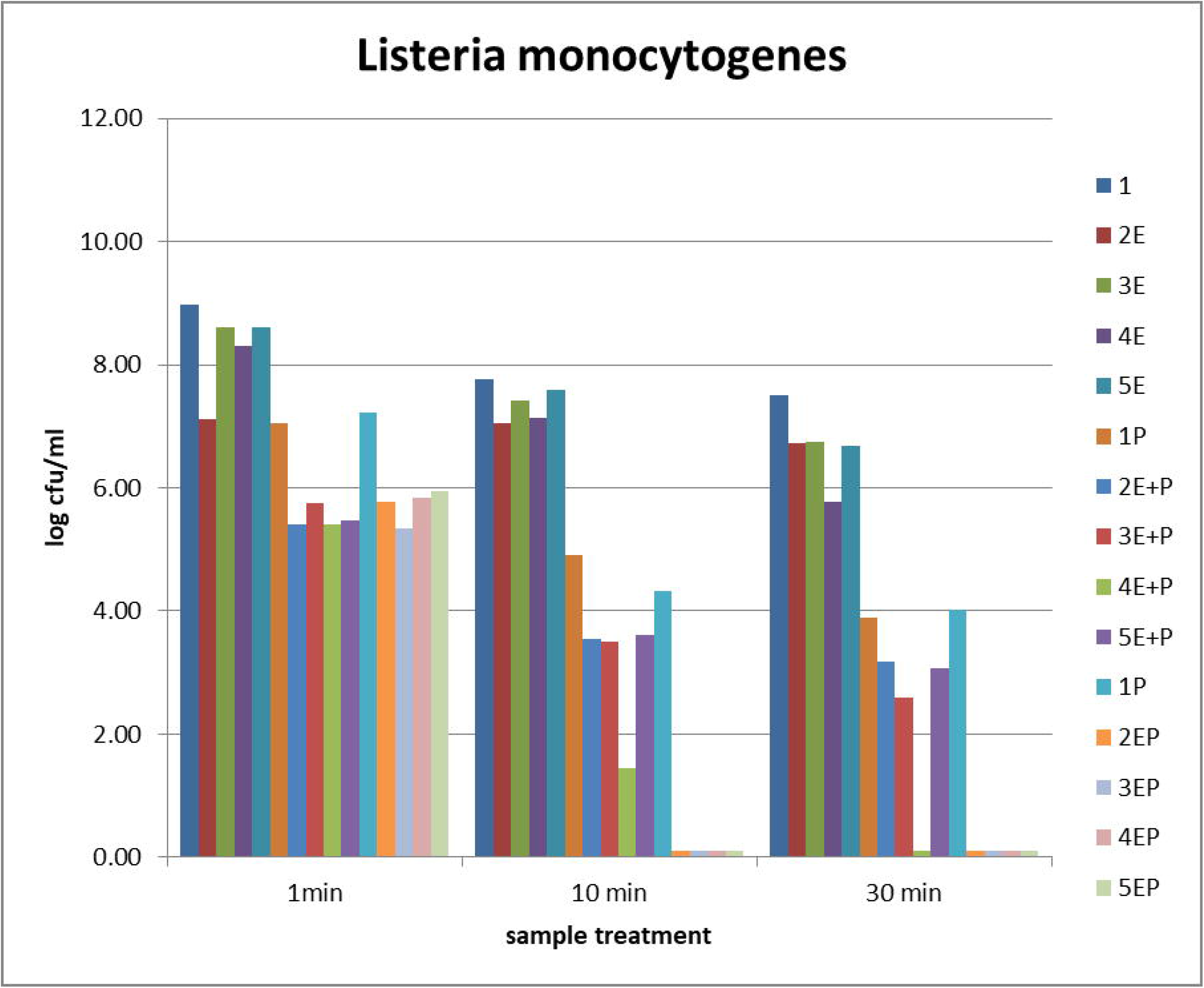
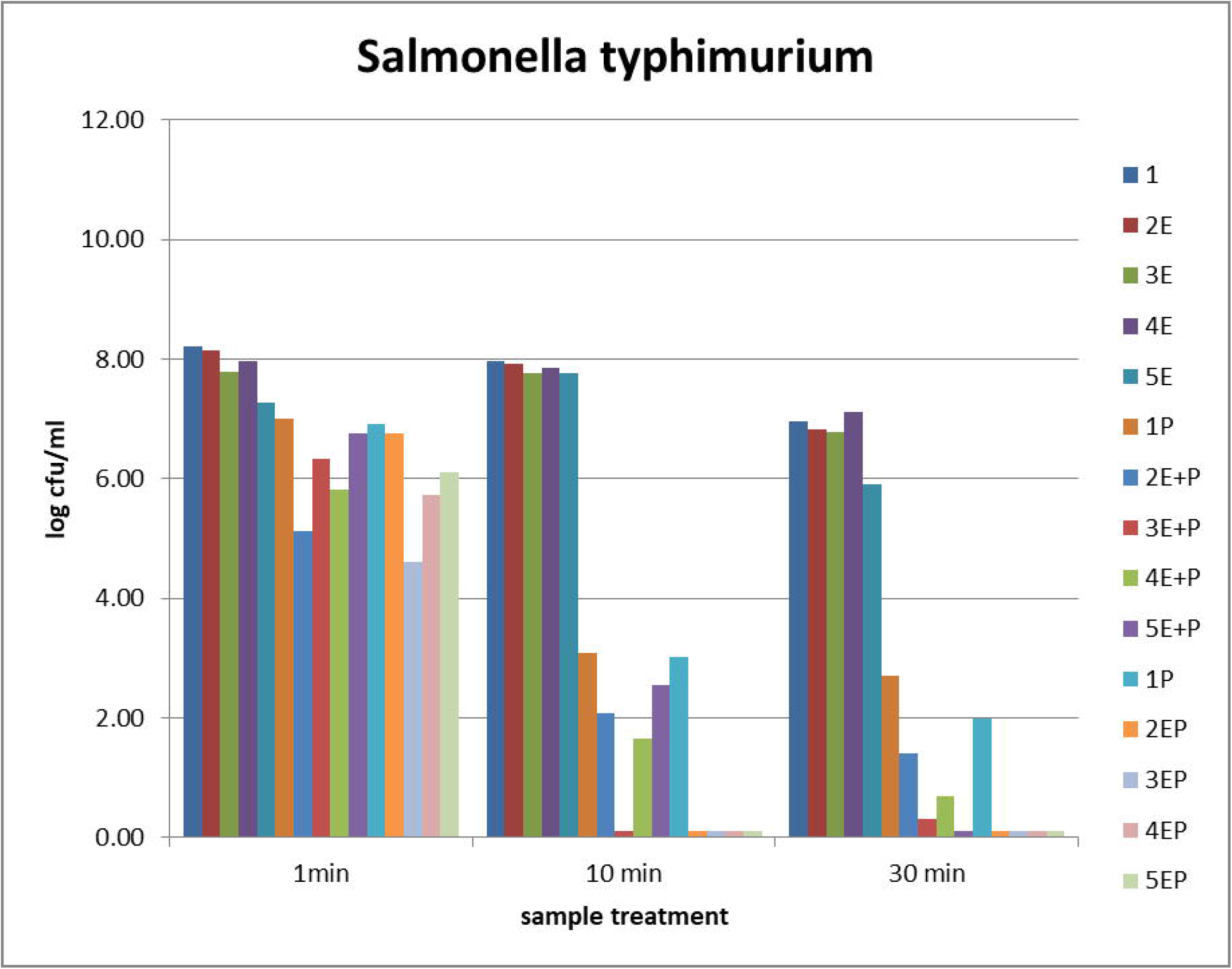
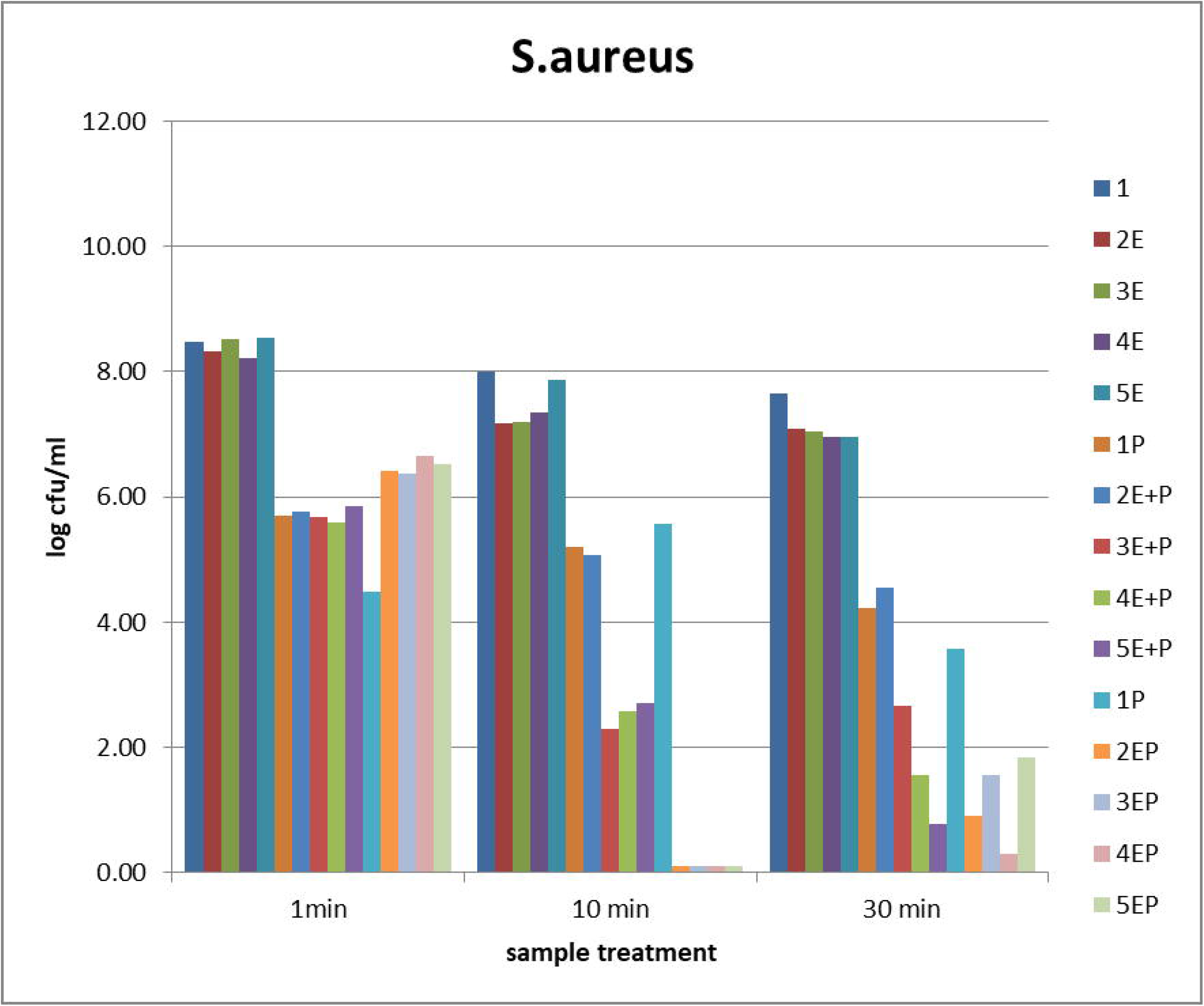

